# Generalization and extinction of learned fear alter primary sensory input to the brain

**DOI:** 10.64898/2026.02.25.708099

**Authors:** Michelle C. Rosenthal, Alper K. Bakir, John P. McGann

## Abstract

Neuroplasticity in sensory systems permits the brain to refine sensory discrimination between threat-predictive and neutral stimuli, but dysfunctional sensory plasticity might underlie maladaptive fear generalization. Using an odor-cued fear conditioning paradigm designed to induce broad fear generalization in a mouse model, we found that odor-evoked synaptic output from olfactory nerve into the brain’s olfactory bulb was greatly increased not only for the original threat-predictive odor but also for novel odors that evoked generalized fear, even under anesthesia. Extinction training in which the threat-predictive odor was presented repeatedly without aversive stimulation reversed the behavioral fear and the increased olfactory nerve output evoked by the threat predictive odor. Extinction training also reversed the generalization of fear and enhanced neurophysiological response to new odors, as did alternative extinction paradigms using novel odorants, thus showing that the output of the olfactory nerve also parallels the generalization of extinction learning. Taken together the increased primary olfactory signaling evoked by fear-evoking odors and the reversal of this increase when the mouse is no longer afraid of an odor suggests that the olfactory nerve plasticity matches the mouse’s perception of threat, even for olfactory stimuli and neuronal populations that have never actually been paired with shock. It is surprising that such beliefs about odor-shock contingencies would manifest as early as the synaptic input from the nose to the brain. This sensory plasticity might contribute to maladaptive generalization of fear such as in post-traumatic syndrome and generalized anxiety disorder.

## Introduction

Fear learning can induce stimulus-specific neuroplasticity in the brain’s sensory systems ^1^. The effects of learning in traditional sensory cortices ^2–5^ and sometimes earlier structures like the thalamus ^6^ have been known for decades. However, it has more recently been appreciated that at least in the olfactory system even the earliest parts of the circuit can be altered by aversive learning ^7–9^, and that changes in odor processing in the brain’s olfactory bulb plays a causal role in shaping fear generalization across odors ^10^. In the olfactory system, the olfactory sensory neurons (OSNs) transduce the odor in the nose and communicate the primary olfactory signal down the olfactory nerve to the brain’s olfactory bulb. In rodent models odor-cued aversive conditioning substantially alters this signal, including odor-specific increases in odor-evoked neurotransmitter release ^11^, odor receptor expression ^9^ and GABA_B_ receptor signaling ^10, 12^. In parallel experiments in humans, these effects are observable in the electroolfactogram recorded intranasally from the olfactory epithelium ^13^, and may drive the conditioning-evoked improvements in olfactory perceptual discrimination reported in participants with normal trait anxiety ^14–17^ but not high trait anxiety ^17^.

Prior studies of learning-induced plasticity in OSNs have emphasized the odor-specificity of the effect to clearly demonstrate that the effects truly reflected learning of the odor-shock contingency as opposed to mere sensitization ^11^. However, the more translationally important question is not whether post-learning OSNs correctly behave as if an odor predicts a shock but whether their plasticity might impact the neural processing of other odors that do not in fact predict shock ^18^. We tested this experimentally using odor-generalized fear learning, where the organism becomes afraid of many related odors, not just the one that predicted a threat. Fear generalization, the physiological or behavioral expression of fear to neutral stimuli that have not been associated with an aversive stimulus, is typically thought to operate in fear networks far removed from the sensory periphery, such as the amygdala, prefrontal cortex, and hippocampus ^19^. However, recent work has demonstrated that conditioning paradigms inducing generalizing fear, where mice become afraid of multiple odors (including novel odors), can cause significant facilitation of neural responses to all those odors at early stages of olfactory processing, including periglomerular interneurons ^20^, mitral cells ^7^ and olfactory cortex ^21^. Given the evidence of odor-specific plasticity in OSNs, we asked whether OSNs might themselves exhibit generalization-related plasticity and thus presumably contribute to or even drive these effects of generalization downstream.

The hypothesized role of OSNs in generalized olfactory fear is unintuitive. Explanations of odor-specific OSN plasticity have emphasized that the OSNs have direct knowledge of the presence of the conditioned stimulus (CS) odor at the time of the shock ^1, 9^. However, during generalizing fear conditioning the subject becomes afraid of odors that are not present and may never have been previously encountered at all. Classical “component-based” models of generalization suggest that this generalization occurs because of overlap in the neural representations of similar sensory stimuli ^22–24^. OSN odor representations can indeed overlap for odors that share chemical features and thus activate some of the same OSN subpopulations ^25, 26^. However, we have previously observed populations of olfactory bulb interneurons whose odor-evoked responses are facilitated by fear generalization from a threat-predictive odor that didn’t excite those interneurons at all, and we have observed OSNs to exhibit a “configural” fear-evoked response modulation, where the same OSN population could be facilitated when driven by a shock-predictive odor and unchanged when driven by a non-shock-predictive odor ^11, 20^. Taken together, these data suggest that generalization of fear across odors has a more complex mechanism than representational overlap.

In learned fear there are two potentially interacting forms of ambiguity: ambiguity about whether a new stimulus predicts a threat like the original stimulus does (i.e. whether to generalize across stimuli) and ambiguity about whether the original threat-predictive stimulus still predicts a threat at a later time (i.e. whether to generalize across time). The latter is usually not conceptualized as generalization *per se*, but rather as “forgetting” (if the ambiguity arises from the mere passage of time) or extinction (if the ambiguity arises from intervening non-reinforced experiences with the original stimulus). Extinction itself has also been interpreted as a generalizable experience, in which the learning of the new stimulus-nonreinforcement contingency might also need to “generalize” to similar stimuli ^27, 28^. We thus tested whether OSN plasticity tracked overall fear across three forms of ambiguity: generalization of learned fear across stimuli, extinction of learned fear across a changed stimulus-threat contingency, and generalization of extinction learning to non-extinguished stimuli. This enables us to test the potential role of sensory plasticity in fear generalization across both stimulus and time ^29^.

Extinction is a form of inhibitory learning ^30^ that is widely used to investigate the behavioral and neural mechanisms of learning and memory ^31, 32^. Previous work in auditory cortex (A1) has shown that extinction learning can reverse learning-induced expansion of the auditory cortical representation of reward-predictive cues ^3^. In the olfactory system, olfactory fear extinction has been shown to reverse learning-induced, stimulus-specific neuroanatomical changes at the primary sensory input to the brain over a long time scale ^33^. However, whether extinction learning reverses the effects of fear learning on sensory neuron physiology remains unknown. It could be ecologically valuable to retain sensitivity gains for a stimulus that was previously highly predictive of a threat ^11, 15^ even if that predictive relationship is no longer true, but such persistence could compete with adaptive responses to current threats. Data from humans suggest that olfactory discrimination performance only partially returns to pre-conditioning baseline after extinction learning ^17^.

Extinction learning offers a translational model to investigate anxiety disorders, including post-traumatic syndrome, and clinical therapies like exposure therapy ^34–36^. Recent work in humans has explored the use of modified extinction or discrimination paradigms using generalization stimuli or alternative approaches with potential therapeutic value ^27, 37–39^. We employed similar variants in mice, including extinction using a panel of disparate odors and extinction-like training using a novel odor, so we could observe the behavioral and neural consequences of these approaches. This may offer translational value to inform clinical exposure therapy treatments, where the original fear-associated cue to be extinguished may not be readily available.

## MATERIALS AND METHODS

### Subjects

Two strains of adult male mice (age mean =20.4 weeks, SD = 3.3) were used in this project, in accordance with protocols approved by the Rutgers University IACUC. Optical neurophysiology experiments were conducted on mice expressing the exocytosis indicator synaptopHluorin (spH), under the control of the olfactory marker protein promoter (*N* = 33) ^11, 40^. These OMP-spH mice, derived from Jackson Labs strain #004946 and reported previously^41–43^, were on an albino C57BL/6 background and were heterozygous for both OMP and spH. Behavioral experiments were conducted on both OMP-spH mice and on wild-type C57BL/6 mice (*N* = 37) from the Jackson Laboratory (Bar Harbor, ME).

### Olfactory Stimuli

A panel of 5 odors was used in this project, consisting of the esters methyl valerate (MV), ethyl valerate (EV), ethyl tiglate (ET), and n-butyl acetate (BA), plus the ketone 2-hexanone (2-Hex). Odors were delivered as described below.

### Behavioral Training

All mice were single housed one week prior to the beginning of experiments. Thirty-three OMP-spH mice were randomly assigned to either the Extinction groups (i.e. the experimental group) or the Procedural “Extinction” group or Never Shocked control group. For a summary of the experimental timeline, please see Fig. 1. All behavioral experimentation took take place in conditioning chambers located inside well-ventilated and sound-attenuated isolation cubicles. For context pre-exposure and fear conditioning, the chamber floors were modular shock floors (16 metal bars controlled by a precision animal shocker) and this was called “Context A”. For extinction training, separate chambers were used, consisting of blue striped walls and white plastic floor for a different visual context from the conditioning chambers, this was “Context B”. All animals underwent context pre-exposure for 15 minutes for two consecutive days. After 48 hours, mice received one session of single cue olfactory fear conditioning consisting of 10 presentations of a 12 second odorant (Methyl Valerate, which was always the shock-predictive conditioned stimulus or CS) at a concentration of 9 arbitrary units (calibrated with a photoionization detector). Each odor presentation co-terminated with a 0.4-mA, 0.5-sec footshock. Trials were separated by a 3.8s −4.8s pseudorandomized intertrial interval. Extinction sessions took place 48 hours after fear conditioning and then every 24 hours for a total of 5 sessions and consisted of the same paradigm as acquisition training except that no shocks were delivered. All behavior was video recorded and freezing scored by Actimetrics Freezeframe. Due to an equipment malfunction, a small minority of odor presentation trials were not adequately video recorded and are not included in the analyses.

**Fig 1.**
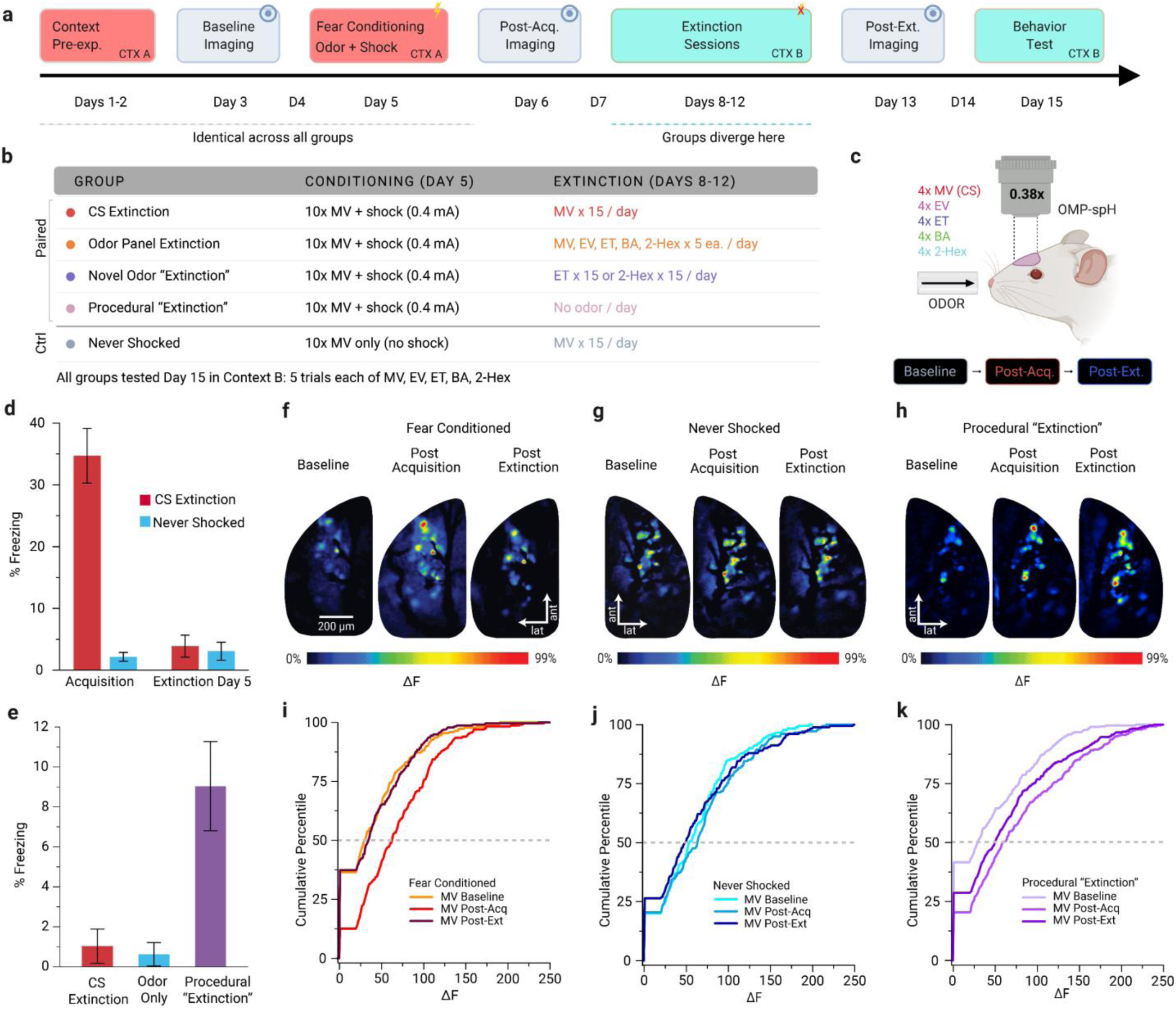
Effects of fear conditioning on response to the CS odor. A) Mice underwent an odor-cued fear conditioning paradigm or control paradigm in context A followed by one of four extinction paradigms in context B. B) Summary of the five experimental groups, clustered as Fear Conditioned/Control based on the fear conditioning or control paradigm. C) Configuration for longitudinal *in vivo* fluorescence imaging of odor-evoked neurotransmitter release from olfactory sensory neurons. D) MV odor-evoked freezing behavior after fear acquisition (left) and after extinction training (right). E) MV odor-evoked freezing behavior after conventional CS extinction, odor only control exposure (Never Shocked), or no-odor (Procedural) sham “extinction”. F-G) Pseudocolor odor-response maps showing the spatial pattern of MV presentation-evoked neurotransmitter release from the olfactory nerve across dorsal olfactory bulb glomeruli before (baseline), after fear learning (post-acquisition), and after extinction learning (post-extinction) for mice in the Fear Conditioned (f), Never Shocked (g), and Procedural “Extinction” only (g) groups. I-k) Cumulative frequency histograms showing the distribution of all MV odor-evoked glomerular response amplitudes in mice from the fear conditioned (i), the Never Shocked control (j), and Procedural “Extinction” (k) groups revealing three different patterns of plasticity. Note the distributional shift from baseline (lightest color) towards larger responses after conditioning (intermediate color) in i and k, followed by a return to the original distribution (darkest color) following MV extinction training in i but not following Procedure Only extinction training in H. Bar graphs in A and B depict the mean ± standard error. Dashed line indicates the median. In all panels odors are abbreviated as MV (methyl valerate, the ester serving as the CS), EV (ethyl valerate, a similar ester), ET (ethyl tiglate, a different ester), BA (n-butyl acetate, a different ester), and 2-Hex (2-hexanone, a ketone).

To test for generalization effects of single-cue olfactory conditioning at the behavioral level, a group of C57BL/6 mice underwent the same fear conditioning and extinction training described above and then received a test comprised of 5 odorant trials of ∼15 seconds each of the CS odorant (MV) and four novel, unexposed odorants (EV, BA, ET, and 2-Hex) varying in similarity to the CS. To investigate whether extinction to the original CS extinguishes generalized fear to novel, unexposed odorants, a second group of C57BL/6 mice received context pre-exposure, fear conditioning and extinction as in Fig. 1 and then received a post extinction test on day 15.

### Longitudinal optical imaging of neural representations of odorants

All surgical procedures for the implantation of chronic cranial windows were previously described ^11^. Briefly, mice were anesthetized with pentobarbital (10 mg/mL, 0.1 mL/10 g, i.p.) and administered additional boosters to maintain anesthetic plane throughout all surgical and imaging procedures. While under anesthesia, body temperature was maintained at 38 ± 0.5°C via a feedback-regulated heating pad. A 0.1% atropine solution was administered (s.c.) to reduce nasal mucous secretions and a 0.25% bupivacaine solution was administered (s.c.) along the incision site as a local anesthetic. The scalp was removed and then fitted with a custom acrylic head cap to permit replicable positioning in the head holder across imaging sessions.

*In vivo* odor-evoked spH signals were acquired using wide-field epifluorescence imaging of the olfactory bulbs, as described previously ^11^. Fluorescence imaging data were collected with a 4× (0.28 NA) objective, and illumination was provided by a 470-nm wavelength bright light-emitting diode with suitable filters. Images were acquired at a pixel resolution of 256×256 at a frame rate of 7 Hz using a monochrome CCD camera. While subjects are secured under the imaging apparatus they were presented with a panel of 5 odors including MV (the CS), the esters EV, BA, and ET, and the ketone 2-Hex. During optical imaging, all odorants were presented at the same concentrations (9 a.u.) used during behavioral training, as calibrated daily via photionization detector (ppbRae, Rae Systems). All odorants were delivered through a manifold located ∼1 cm in front of the mouse’s nose via a custom vapor dilution olfactometer operated through MATLAB-based software. Each odorant stimulus was presented in a block of 4 individual trials separated by 60-sec ITIs, with each individual trial consisting of 112 acquired frames that were comprised of a 4-sec pre-odorant baseline, 6-sec odorant presentation, and 6-sec post-odorant recovery period. To improve signal to noise ratio, the 4 trials for each odorant block were averaged. Two blocks of blank (no odorant) trials were presented at the beginning and end of each imaging session, and were averaged together and subtracted from odorant trials to correct for photobleaching.

### Data Analysis

For behavioral experiments, freezing was defined as a lack of movement except for respiration during odorant trials and the proportion of time spent freezing on each trial was computed using FreezeFrame 4. Data was exported to Excel, SPSS, and Origin Pro for statistical analysis (see below).

Fluorescence imaging data were quantified and analyzed as previously reported ^11^. Glomerular regions of interest (ROIs) were identified and hand-selected on response maps consisting of blank-subtracted average maps, for each odorant and each concentration. To quantify the peak odorant-evoked change in fluorescence (ΔF), spH signals for each trace corresponding to a glomerular ROI were determined by subtracting 1 sec of baseline frames (acquired during the pre-odorant baseline) from 1 sec of frames centered around the peak trace inflection after odorant onset. Because spH provides an integrative signal of exocytosis over time ^40^, the response peak magnitudes typically occur towards the end of the 6-sec odorant presentation; frames 63-76 were thus used for this subtraction. All data were high-pass filtered with a Gaussian filter through software written in MATLAB and exported to Excel, SPSS, and OriginPro for statistical analysis (see below). ROIs were operationally defined as responding to an odor if their average odor-evoked increase in fluorescence across a block of odor presentations was at least 20 fluorescence units, which corresponds to an approximately five-fold increase above root-mean-square noise in our spH fluorescence signals.

To compare changes in overall response distributions between training sessions, nonparametric two-sample Kolmogorov-Smirnov tests were used. To determine changes in central tendencies of groups based on means from individual subjects, parametric and non-parametric tests were used as appropriate. An α level of 0.05 were used to accept statistical significance and was Bonferroni corrected for multiple comparisons within analyses. To examine the effects of extinction training on reversing the effects of aversive-learning, mice that did not show an initial change from baseline to the post-conditioning imaging session for a given stimulus were excluded for extinction-related reversal analyses involving that stimulus, using the following criterion: the ratio of mean ΔF post-acquisition/ mean ΔF baseline > 1. This excluded 1 mouse from the CS Extinction group, 2 mice from the Novel Odor “Extinction” group, 1 mouse from the Odor Panel extinction group, and 2 mice from the Procedural “Extinction” group.

## RESULTS

### Experimental Paradigm

We used an odor-cued fear conditioning and extinction paradigm for mice that can be completed in 8 days of behavioral training, thus permitting all behavioral training and three imaging sessions to be performed within 15 days on each mouse. This reduces the potentially confounding effects of structural plasticity in the olfactory epithelium, which has been observed after roughly three weeks ^9^. The experimental sequence is illustrated in **Fig. 1a & b**. During each imaging session we used optical neurophysiological methods ^40^ to record neurotransmitter release *in vivo* from populations of axon terminals of mature olfactory sensory neurons (OSNs) using OMP-spH gene-targeted mice. OSNs are the primary sensory neurons in the olfactory system, transducing the odorant in the olfactory epithelium and projecting their axons to the brain’s olfactory bulb, where they sort by odor receptor type into a sheet of glomeruli across the bulb’s surface. Odor presentation under the microscope (**Fig. 1c**) evokes increases in fluorescence in an odor-specific subset of glomeruli, which indicates the primary sensory representation of the peripheral odor stimulus at the input to the brain. Each individual mouse was imaged at three time-points: before olfactory fear conditioning, after olfactory fear conditioning (or odor exposure controls in the Never Shocked group), and after extinction training (or Procedural “Extinction” control exposure) as listed in **Fig.1b**. Freezing behavior during extinction training was quantified to confirm learning. Animals were awake during training but were lightly anesthetized during optical imaging to isolate “bottom-up” signaling in the olfactory nerve from any “top-down” factors like changes in respiration in expectation of shock ^11^.

### Odor-evoked OSN output is reversibly increased for a threat predictive odor

As shown in **Fig. 1a**, Mice were exposed to the fear conditioning chamber (context A) for two pre-exposure sessions, then received a single fear conditioning session of ten trials in which the fruity odor methyl valerate (MV), an ester, was blown into the chamber for 12s and paired with an aversive electrical footshock (Fear Conditioned groups). Mice in the Never Shocked control group received the same conditioning but with the shocker switched off. The fear conditioning session was followed by five additional daily sessions of extinction training in a different chamber (context B), in which subsets of mice received conventional CS Extinction (15 daily presentations of MV without shock for a total of 75 extinction trials) or Procedural “Extinction” (same as CS Extinction but with the odor turned off). The Never Shocked group also underwent the CS Extinction paradigm to control for the effects of the additional odor exposure, but these are referred to as the Never Shocked “Extinction” group because there was no fear to extinguish in these mice. Mice then underwent a final behavioral testing session at the end of the experiment in which multiple test odors were presented in context B. OSN synaptic output from the olfactory nerve into olfactory bulb glomeruli was assessed via optical neurophysiology during a pre-conditioning baseline session, again after fear conditioning, and again after extinction. Additional groups that instead received Odor Panel Extinction or Novel Odor “Extinction” paradigms (**Fig. 1b**) are discussed below.

The behavioral training successfully induced conditioned fear of the CS odor (always MV), followed by robust extinction of odor-evoked conditioned fear. There was a statistically significant interaction between training session and experimental group on odor-evoked freezing behavior (*F*_1,31_ = 85.15, *p* < 0.001, η_p_^2^ = 0.73). At the end of the fear conditioning session (**Fig. 1d**), we observed robust odor-evoked freezing in mice in the odor-shock Fear Conditioned groups compared to mice in the Never Shocked group (*F*_1,31_ = 80.71, *p* < 0.001, η_p_^2^ = 0.72). Five sessions of extinction with the CS odor was enough to significantly reduce odor-evoked freezing in the Fear Conditioned group (**Fig. 1d**) compared with their freezing during acquisition (*p* < 0.001), making their levels of freezing no different than animals in the Never Shocked group (*p* = 0.879). Levels of odor-evoked freezing for mice in the control Never Shocked group did not differ from the acquisition training session to day 5 of extinction training (**Fig. 1d**; *p* = 0.501). During the final behavioral test (**Fig. 1e**), there was a significant effect of extinction group on CS-evoked freezing F_2,39_ = 14.91, *p* < 0.001, η_p_^2^ = .43). Mice in the Procedural “Extinction” group, which underwent the extinction paradigm without any odor presentations (*N* = 10, *M* = 9.03, *SD =* 7.06) exhibited significantly more freezing than animals in the CS Extinction group (*N* = 13, *M* = 1.02, *SD* = 3.09, *p* <0.001) and Never Shocked group (*N* = 19, *M* = 0.62, *SD* = 2.54, *p* <0.001), demonstrating some retention of the conditioned fear response even when tested in a different context after days of sham extinction sessions without odor presentations. CS-evoked freezing for mice in the MV extinction group was not statistically different from freezing for mice in the Never Shocked group (*p* = 1.000) confirming the efficacy of the conventional extinction paradigm.

The first hypothesis was that CS odor-evoked OSN output would be facilitated after odor-cued fear conditioning, as previously reported ^11^, and then returned to baseline following extinction learning. We thus analyzed longitudinal changes in MV-evoked OSN output from the mice in the MV Extinction group at baseline, then again following acquisition of MV-cued fear, and then again following extinction of MV-cued fear. As shown in **Fig. 1F and I**, these mice (*N* = 5) displayed a robust increase in CS-evoked OSN synaptic output following fear conditioning compared to the pre-conditioning baseline (Kolmogorov-Smirnov test; *N* = 452 glomeruli, *D = 2.89,* p <0.001). Subsequent CS Extinction training then decreased the MV-evoked OSN output compared to the post acquisition imaging session (*N* = 8, 645 glomeruli, *D* = 2.96 *p* <0.001), returning the CS-evoked OSN output to be no different from its original distribution of response amplitudes (*N* = 645 glomeruli, *D* = 0.658, *p* = 0.780). This pattern shows for the first time that the OSN signals (analogous to a photoreceptor or inner hair cell) both increase and decrease to track the expected outcome (shock or no shock) corresponding to the odorant.

In contrast, MV-evoked OSN output did not change from baseline to post acquisition for animals in the Never Shocked control group (**Fig. 1g & j**; *N* = 645 glomeruli, D = 0.98, *p* = 0.288), and modestly decreased after the extended MV exposure in the subsequent “extinction” paradigm (*N* = 645 glomeruli; *D* = 1.80, *p* = 0.003 vs. post-acquisition; *D* = 1.79, *p* = 0.003 vs. baseline). This confirms that the mere exposure to the CS odor was insufficient to increase OSN output over the course of the conditioning paradigm and could somewhat decrease it after many exposures ^42^. Mice in the Procedural “Extinction” control group (*N* = 5), exhibited the expected increase in CS-evoked OSN output after fear acquisition (**Fig. 1h & k**) compared to the pre-conditioning baseline (*N* = 622 glomeruli, D = 2.93, *p* < 0.001), but the five days of procedural “extinction” without odors induced only a slight decrease in CS-evoked OSN output compared to their post-acquisition imaging session (*N* = 622 glomeruli, *D* = 1.36, *p* <0.049), leaving it significantly elevated compared to the pre-conditioning baseline (*N* = 622 glomeruli, *D* = 1.84, *p* = 0.002). This demonstrates that extensive exposure to the extinction context in the absence of explicit odor presentation did have a small effect on OSN output, but not nearly as large as when the CS odor was presented (**Fig. 1i**).

### Olfactory aversive conditioning induced generalization of fear to diverse odors

The degree of fear generalization across odors is determined by the details of the fear conditioning paradigm. We have previously used paradigms intended to produce odor-specific fear ^11^, a generalization gradient across similar odors ^10, 44^, or widely generalized fear across disparate odors ^20^. Here we used a fear conditioning paradigm that promoted widely generalized fear so that we could experiment with the generalization of subsequent extinction training. When a test panel composed of the CS odor (MV) and four novel odors was presented in the novel test context (context B) three days after fear conditioning, mice displayed significant odor-evoked freezing to all five odors (**Fig. 2g**; Wilcoxon signed-rank tests, BA: p < 0.002, other odors: p < 0.001). This confirms that (as intended) mice generalized their fear comparably from methyl valerate (MV) to the very similar ester ethyl valerate (EV), to the somewhat similar esters n-butyl acetate (BA) and ethyl tiglate (ET), and to the quite different smelling ketone 2-hexanone (2-Hex).

**Fig. 2.**
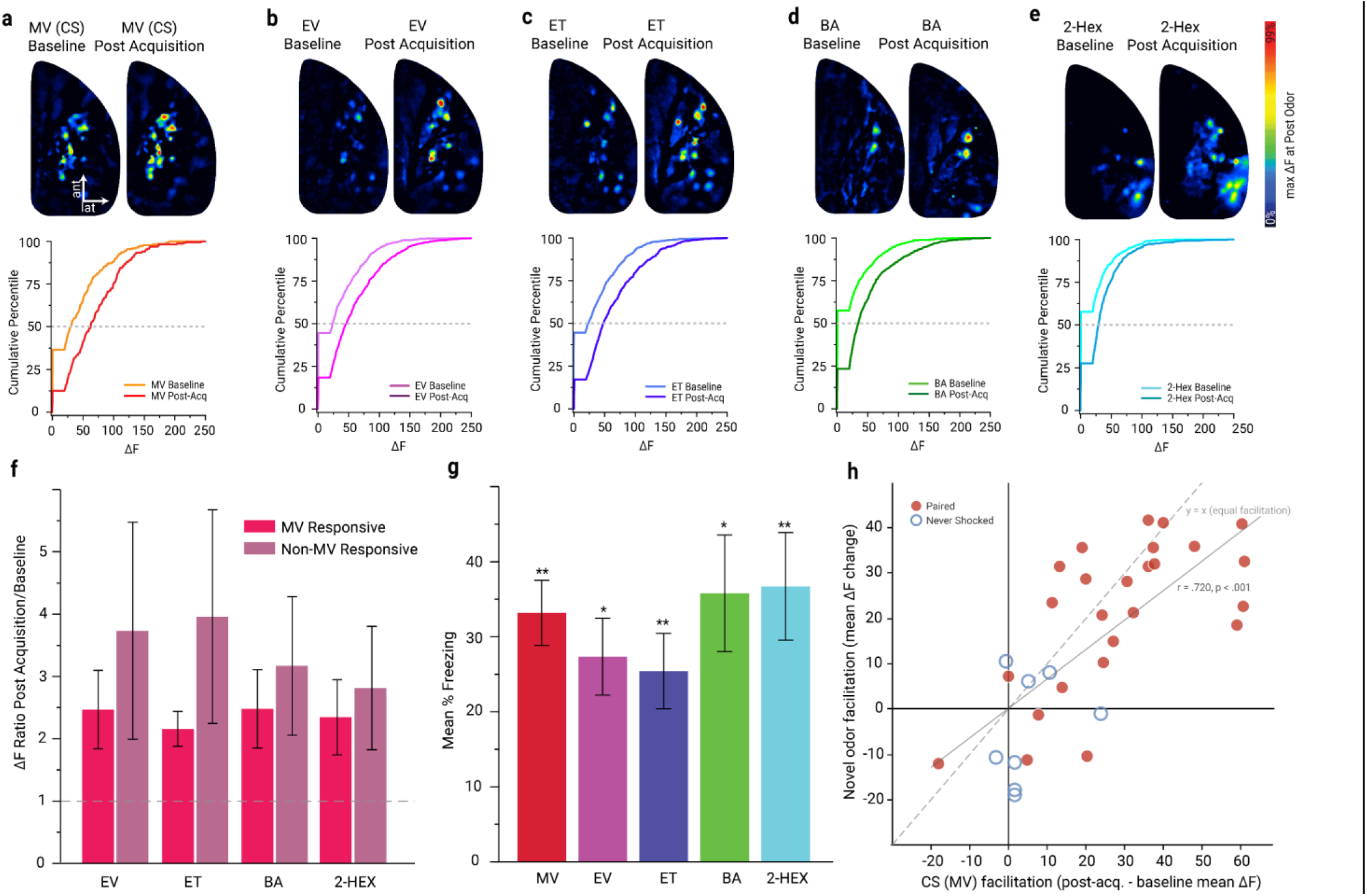
Generalization of fear learning and OSN neuroplasticity with novel odors. **(A-E)** Top: Pseudocolored odor response maps from a representative mouse from each group showing change from before conditioning (left) to after conditioning (right). Bottom: Cumulative frequency histograms of odor-evoked glomerular responses demonstrating olfactory aversive conditioning increases the number of responsive glomeruli and the size of the responses four all odors in the panel for mice in the Fear Conditioned groups. (**F)** Mean ratio of odor-evoked glomerular response amplitudes ± SEM for MV-responsive and non-responsive glomeruli demonstrating that the learning-induced change was at least as big in glomeruli that didn’t respond to the CS (purple) as in those that did respond to the CS (red). Dashed line represents no change from baseline to post acquisition in odor-evoked spH responses. (**G)** Mean odor-evoked freezing behavior during first 2 trials of extinction day 1 (3 days after acquisition) for each odor in the panel. **(H)** Correlation between degree of change in MV-evoked signals and change in novel odor-evoked signals, pooling across odor-shock paired Fear Conditioned (closed circles) and Never Shocked groups (open circles). Double asterisk represents (p< .001), single (p<.05). In all panels odors are abbreviated as MV (methyl valerate, the ester serving as the CS), EV (ethyl valerate, a similar ester), ET (ethyl tiglate, a different ester), BA (n-butyl acetate, a different ester), and 2-Hex (2-hexanone, a ketone).

### Generalization of learned fear was accompanied by increases in OSN output, even for OSNs that don’t respond to the CS

The fear generalization paradigm allowed us to ask whether fear conditioning-induced changes in the output of OSNs reflect the actual history of odor-shock pairings or instead follows the mouse’s generalized expectation of odor-signaled threat even for novel odors. As displayed in **Fig. 2a-e**, mice in groups receiving paired odor-shock training in the Fear Conditioned group exhibited large increases in odor-evoked OSN output after MV-cued fear learning, not just to MV (**Fig. 2a**) but to all odors tested (**Fig. 2b-e**; K-S tests; N ranged from 1290 to 1859 glomeruli depending on odor; D ranged from 5.49 to 6.72; all p < 0.001). The effects were comparable in size across odorants, regardless of similarity to the CS and despite their novelty to the mouse. Behavioral fear generalization to new odors was thus accompanied by OSN neuroplasticity for those new odors as well.

A classical model for generalization across stimuli posits that similar stimuli share elements in common, and those shared elements convey the learned response across stimuli in proportion to the degree of overlap ^45^. In the early olfactory system the shared elements among odor representations are individually observable, where a given odorant molecule binds to a subset of odor receptors and drives OSN output into a corresponding subset of olfactory bulb glomeruli. Two chemically similar odorants will both bind to some of the same receptors and consequently excite overlapping sets of olfactory bulb glomeruli. We thus asked whether the increased OSN output in response novel odors after generalizing fear conditioning could result from overlapping sets of glomeruli between each test odor and the CS odor. Remarkably, the subset of glomeruli that responded only to the novel test odorant and not to the CS exhibited significantly enhanced responses following fear conditioning with the CS odor (**Fig. 2F**; one-sample *t-*test vs. baseline*, p* values *<*0.05 across all odors), and in fact the size of the increase was if anything slightly larger for the glomeruli that didn’t respond to the CS than for those that did for all four test odors (**Fig. 2F**, purple bars vs red bars). This result, combined with the comparable freezing across odors regardless of similarity to the CS (**Fig. 2g**), suggests that the generalization of learned fear across novel odors is driven by the mouse’s inference about odor meaning rather than peripheral overlap in odor response.

### Effects of conventional extinction training on odor-evoked OSN output across odors

Given that fear conditioning with a single CS odor evoked generalized fear and corresponding facilitation of OSN output across disparate odors, we next asked whether CS extinction training would also reverse the increase across disparate odors. As described earlier, conventional extinction using the CS itself almost perfectly reversed the facilitated neural response to the CS (**Figs. 1i & 3a, left**). However, conventional extinction with the CS odor did *not* fully reverse the effects of fear generalization. As shown in **Fig. 3a**, for all four novel odors the distribution of OSN outputs after conventional CS extinction (darkest lines) was reverted only partway back towards baseline (lightest lines) from the post-conditioning distributions (medium lines). The partial reversion was significantly different from the post-conditioning distribution for all four odors (K-S tests; *N* range: 100-306 glomeruli; *D* range: 1.83-3.30; BA *p* = 0.003, other *p*’s < 0.001), but it remained significantly elevated for BA (*N* = 301 glomeruli from 5 mice, *D* = 2.39, *p* < 0.001) and ET (*N* = 246 glomeruli from 4 mice, *D* = 1.47, *p* = 0.027). The reversal of the fear-induced OSN facilitation for the novel test odors after CS extinction largely corresponded to the freezing behavior, which was reduced to near zero for every odor following this conventional extinction paradigm (**Fig. 3e**), with the exception of modest but significant freezing evoked by 2-Hex (Wilcoxon; *N* = 13, *Z* = 2.02, *p* = 0.043). Between the continued freezing to 2-Hex (the most dissimilar odor to the CS) and the partial retention of the facilitated neurophysiological response to all of the test odors but not the CS itself (**Fig. 3a**), we interpret this pattern of results as evidence that the generalization of fear extinction was narrower than the generalization of fear learning.

**Fig. 3.**
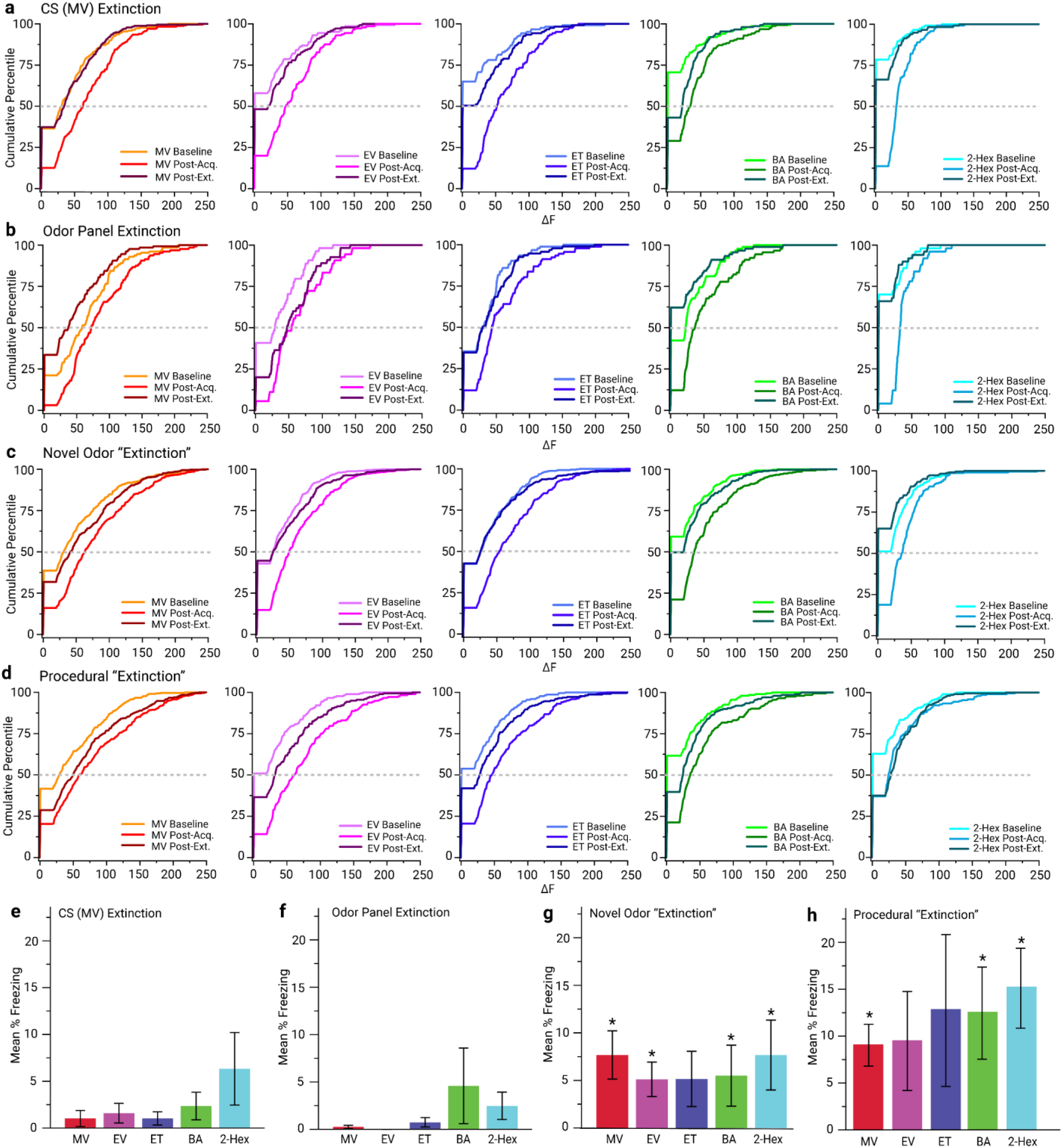
Effects of different extinction paradigms on olfactory neurophysiology and behavior. **(A-D)** Cumulative frequency histograms of odor-evoked responses demonstrating effects of olfactory aversive conditioning and conventional extinction training by comparing baseline (lightest color), post-acquisition (medium shade), and post-extinction (darkest shade) for the CS odor (left) and other odors. Data from the MV Extinction group (a), Procedural “Extinction” group (b), Odor Panel Extinction group (c), and Novel Odor “Extinction” group (d) are shown. Dashed lines indicate the median. **(E-H)** Mean percent freezing evoked by each test odor during the behavioral test session in context B mice receiving conventional MV extinction training (e), odor panel extinction (f), novel odor “extinction” (g), or procedural “extinction” (h). Asterisk indicates *p* < 0.05. In all panels odors are abbreviated as MV (methyl valerate, the ester serving as the CS), EV (ethyl valerate, a similar ester), ET (ethyl tiglate, a different ester), BA (n-butyl acetate, a different ester), and 2-Hex (2-hexanone, a ketone).

### Odor panel extinction training

Given the incomplete effect of the conventional extinction paradigm on novel odors, we also explored the behavioral and neurophysiological effects of an alternative paradigm in which we replaced the conventional CS extinction training (75 presentations of MV over 5 days) with an Odor Panel Extinction paradigm in which each of the odors was presented 5 times per day (interleaved) for five days. This reduced the total number of true extinction trials of the actual CS (MV) from 75 to 25, while adding 25 presentations each of the novel odors EV, ET, BA, and 2-Hex. We hypothesized that this paradigm might “refine” the learned fear, demonstrating that the novel test odors do not predict a shock while leaving some of the fear associated with the actual CS intact.

Surprisingly, this Odor Panel Extinction not only didn’t leave some of the CS-associated fear intact, it actually had a much larger effect on the response to the CS than conventional extinction did despite the greatly reduced number of CS presentations. As expected, fear learning greatly increased the odor-evoked glomerular response to MV and all four novel odors as above (**Fig. 2**) in this subset of mice. However, subsequent Odor Panel Extinction training so strongly reduced the fear learning-facilitated OSN responses to MV (*N* = 256 ROIs, *D* = 3.00, *p* < 0.001) that it drove the post-extinction OSN response (**Fig. 3c, darkest line**) significantly below baseline (**Fig. 3c, lightest line**; *p* < 0.001). This is consistent with the behavioral data (**Fig. 3f**) showing that MV-evoked freezing was completely abolished following odor panel extinction.

All four novel odors exhibited significant returns toward baseline (K-S test; *N* range: 180-812 glomeruli; *D* range: 3.35-4.89; all *p*’s < 0.001), with ET and 2-Hex returning to their baseline distribution following odor panel extinction and BA overshooting to become less than baseline. Note that the EV data is limited to two mice in this group due to technical problems during data collection. None of these odors evoked significant freezing after Odor Panel Extinction (**Fig. 3f**; one-sample Wilcoxon signed-rank *N* = 9, all *p’*s > 0.05). This was the only extinction paradigm in which mice that had received aversive learning displayed no statistically significant levels of freezing to any of the odors in the panel. Based purely on the number of CS presentations, odor panel extinction (15 CS presentations) was more notably more effective at extinguishing freezing than conventional CS extinction (75 CS presentations), in which mice still averaged 37% freezing over extinction trials 11-15.

### Novel Odor “Extinction”

Because the odor panel paradigm was so effective at reversing the behavioral and neurophysiological effects of fear learning despite its reduced number of CS presentations, we hypothesized that a paradigm presenting *only* novel odors might be sufficient to reverse some of the effects of odor-cued fear conditioning. In this paradigm we presented 15 daily trials of an arbitrary odor from the test panel for five days instead of conventional CS extinction training. For 9 mice we used ET and for 14 mice we used 2-Hex, then we pooled these data together afterwards. These animals received zero “true extinction” trials in which the CS is presented without shock (and we thus put quotes around “extinction” for this paradigm), and any change in the behavioral or neurophysiological response to MV can thus be considered generalization of extinction.

As expected, fear learning greatly increased the odor-evoked glomerular response to MV and all four novel odors as above (**Fig. 2**) in this subset of mice. Surprisingly, “extinction” training with a novel odor strongly reversed this fear learning induced enhancement to MV (**Fig. 3d**), significantly reducing MV-evoked OSN output relative to the post-fear conditioning state (*N* = 859 glomeruli from 9 mice, *D* = 3.22, *p* <0.001) and leaving the response slightly elevated but not significantly different from the pre-conditioning baseline (*N* = 860 glomeruli, *D* = 1.42, *p* = 0.35). Freezing evoked by MV was substantially less than exhibited prior to the “extinction” training (**Fig. 2d**), though it remained significant (**Fig. 3g**). The novel odor “extinction” paradigm likewise reversed the facilitated OSN responses induced by MV-cued fear conditioning for all four test odors (**Fig. 3d**; K-S tests; *N* range: 446-816 glomeruli; *D* range: 3.65-4.77; all *p*’s less than 0.001), returning to or slightly below baseline responses. As for MV, freezing evoked by the test odors was substantially less than exhibited prior to “extinction” training (**Fig. 2g**), though it remained significant (**Fig. 3g**) for all odors except ET (Wilcoxon signed-rank; *Z* = 1.83, *p* = 0.068). This demonstrates that extinction-like experience with a different odor can generalize its behavioral and neurophysiological effects to the original CS and other novel odors, though not as effectively as exposure to the CS itself or a panel including the CS.

### Generalization of Procedural “Extinction” to OSN physiology

In the highly generalizing fear conditioning paradigm employed here, mice generalized their learned fear to novel odors (**Fig. 2**), with effects even in OSN populations that were not engaged by the CS itself (**Fig. 2f**), and also generalized their learned extinction such that exposure to completely novel odors reduced both the fear and the facilitated OSN output evoked by the CS itself (**Fig. 3c, d, f, and g**). Since the presentation of the CS is thus not necessary to reverse the effects of conditioning, we asked whether any odor need be presented at all or whether the context of being handled and placed in the apparatus could be sufficient to reverse some of the effects of odor-cued fear conditioning. We thus also performed a no-odor Procedural “Extinction” paradigm, in which fear conditioned mice underwent identical procedures to the CS Extinction group but without any odor presentations (i.e. for 5 days in Context B). We place “extinction” in quotes because this paradigm does not include any unreinforced CS presentations.

Because the Procedural “Extinction” group does not experience odors during extinction, we compare their freezing behavior to other groups. Procedural “Extinction” partially extinguished CS-evoked fear, such that mice dropped from MV-evoked freezing about 32% of the time (**Fig. 2g**) after conditioning to MV-evoked freezing about 8% of the time after Procedural “Extinction” (**Fig. 3g**), though still more than the 1% CS-evoked freezing observed after conventional CS Extinction (**Fig. 3e**). Mice that underwent Procedural “Extinction” also exhibited less freezing in response to EV, BA, and 2-Hex (**Fig. 3h vs. Fig. 2g**), though significant freezing remained to all odors but ET (Wilcoxon; *p* < 0.05).

As expected, fear learning greatly increased the odor-evoked glomerular response to MV and all four novel odors as above (**Fig. 2**) in this subset of mice. Following the Procedural “Extinction” paradigm, in which the mice were placed in the extinction context for five daily sessions but no odors were presented, the MV-evoked OSN outputs were modestly but significantly reduced relative to their post-conditioning state (**Fig 3b**; *N* = 622 glomeruli from 6 mice; *D* = 1.63; *p* = .049). Procedural “Extinction” also reduced glomerular responding for all four novel odors in the panel (**Fig. 3b**; *N* range=358-448 glomeruli; *D* range = 1.74-2.50; 2-Hex p=0.004, all other *p*’s <0.001). However, the distribution of glomerular responses evoked by all odors remained elevated compared to baseline (**Fig. 3b**), indicating that the response magnitude for all odors tested remained enhanced compared to baseline (all *p* <0.05). Overall, Procedural “Extinction” had the smallest effect of the 4 extinction paradigms, but it provides some evidence that even mere exposure to the experimental procedure or perhaps the mere passage of time can cause early olfactory processing to partially revert to its pre-conditioning state.

### Aversive learning and extinction learning alter neural representations of odors

Each glomerulus in the olfactory bulb corresponds to a particular type of odor receptor in the nose. Odors are thus first represented in the brain by the set of glomeruli that receive OSN input. We abstracted each pattern of odor-evoked activity across the dorsal aspect of the two olfactory bulbs as a vector with elements corresponding to the set of all glomeruli that responded to any odor whose activity changes over the duration of an odor presentation. This is illustrated in the heat maps in **Fig. 4a**, which each show the timecourse of the fluorescence signal in each glomerulus, where time is depicted from left to right and each glomerulus is one row. By comparing these heat maps from before (PRE) to after (POST) conditioning, it is easy to see which glomerular responses became larger or were added to the vector following odor-cued conditioning (**Fig. 4a**, top two rows) or following odor-only control exposure (**Fig. 4a**, third and fourth rows).

**Fig 4.**
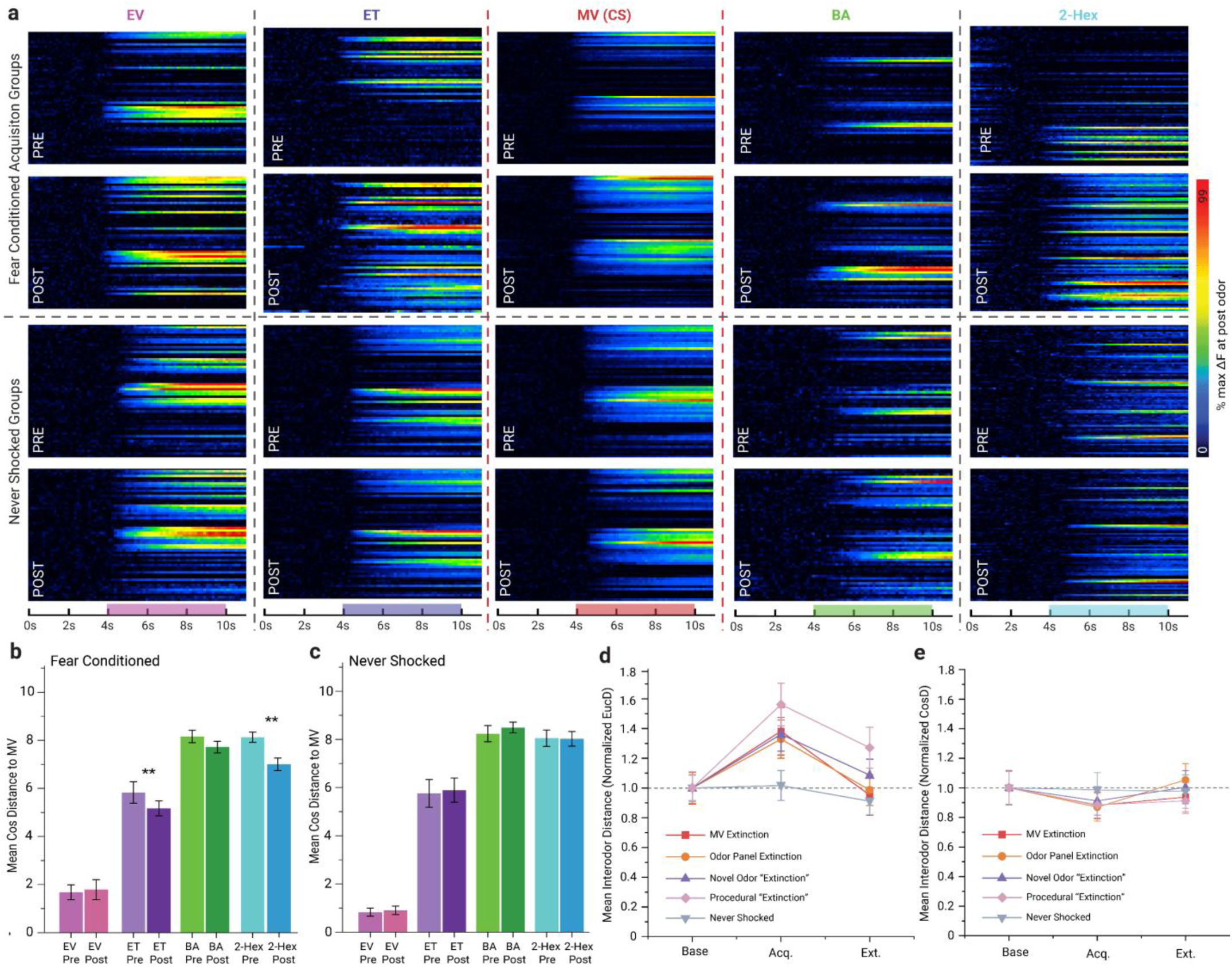
Fear and extinction learning alter primary odor representations. (**A)** Pseudocolored heat maps of odor-evoked activity over time within an odor presentation across all glomeruli in representative mice. Each map represents a single odor presentation in a single mouse, where each row is the response of an individual responsive glomerulus before, during, and just after the odor presentation. Scale bar displays the timing of odor presentation within a trial, in seconds. Responses are scaled to 99% of the peak response post-conditioning. The top two rows are all from one mouse in the Fear Conditioned group before (top row) and after (second row) fear conditioning. The third and fourth row are all from one mouse in the Never Shocked control group before (third row) and after (fourth row) odor exposure. Note that each heat map defines a time-varying vector in a space defined by the responsive glomeruli that represents the identity of the odor. (**B-C**) Mean cosine (amplitude-independent) distances between the neural representation of each test odor and the neural representation of MV (the CS) before (Pre) and after (Post) fear conditioning. Note that the more distant (i.e. less similar) odors become more like MV pooled across the fear conditioned groups (b) but not in the odor only (Never Shocked) control group (c). **D)** Average Euclidean (amplitude-dependent) distance between MV and the four other test odors before learning (base), after fear acquisition (Acq), and after extinction or control exposures (Ext.) showing the increased distances following fear conditioning but not odor alone (green) that is then restored to baseline following extinction. **E**) Same as D but with cosine (amplitude-independent) distance shows that the modest reduction in distance caused by fear learning is eliminated following fear extinction. Error bars represent 1 SEM throughout.

An advantage of these vectoral representations is that they enable the quantitative measurement of odor dissimilarity as the distance between any pair of odor responses in an individual mouse in vector space, either as the response magnitude-independent cosine distance between the odor representations (the angle between the two vectors) or as the response magnitude-dependent Euclidean distance between the odor representations (the absolute distance between the tips of the vectors). **Fig. 4b** shows the cosine distance between each test odor and the CS odor (MV) before and after fear conditioning for the Fear Conditioned group. Note that before conditioning the cosine distance captures the relative dissimilarity across odors, with EV being very similar to MV (short distance), BA and 2-Hex both quite dissimilar to MV (long distance), and ET being in between in similarity (medium distance). Following MV-cued conditioning, the representations of ET and 2-Hex became significantly less different from MV (ET: *t*_25_ = −5.77, *p* < 0.001, Cohen’s *d* = −1.1; 2-Hex: *t*_23_ = −3.68, *p* < 0.001, *d* = 0.75), with large statistical effect sizes though modest changes relative to the already large dissimilarities between them. The Never Shocked group, by comparison, showed no change in cosine odor dissimilarity (**Fig. 4c**, all *p*’s >0.26). The distances can alternately be quantified by Euclidean distance which, unsurprisingly given the amplitude increases shown above, shows large increases in dissimilarity from MV for all four odors (t-test; N=26, *t* range: 3.73-6.65, *d* range: 0.73-1.3, all *p*’s < 0.001).

We then turned to the extinction manipulation to see how representations changed, though that requires parsing the data into smaller groups. **Fig. 4d** shows the relative change in the average Euclidean distance between the CS odor and the other four odors for each extinction group. The larger OSN outputs after conditioning result in much larger dissimilarities between odors for all four extinction groups but no change for the Never Shocked group. Following extinction training, the average Euclidean distance between MV and the test odors decreased for all fear conditioned groups (**Fig. 4d**). The average cosine dissimilarity between MV and the test odors decreased for all fear conditioned groups (**Fig. 4e**) then returned to baseline after extinction, while the Never Shocked group (green) didn’t change. Taken together, these data demonstrate that odor representations at the output of the olfactory nerve not only capture the chemical identity of the odor, but also reversibly change to become both further apart in an absolute sense and more similar in a relative sense when odors are believed to predict the same bad outcome.

### Relationship between OSN neurophysiology and behavioral outcomes under different extinction conditions

As shown in Fig. 5, we observed the different extinction paradigms to produce generally corresponding outcomes between OSN physiology and observed fear behavior in response to the CS odor. The paradigms that produced the least net increase in OSN response amplitudes between the beginning and end of the experiment (CS Extinction, Odor Panel Extinction, and Never Shocked Control groups) also resulted in mice that did not freeze in response to any odor. The two paradigms that left residual increases in OSN response amplitudes following fear conditioning and extinction (Novel Odor “Extinction” and Procedural “Extinction”) resulted in mice that exhibited modest amounts of freezing. Fear conditioning alone, prior to any form of extinction, produced both large increases in OSN output and the largest amount of freezing (Fig. 1). This is consistent with recent results demonstrating that alterations in olfactory bulb GABA_B_ receptor signaling (which presynaptically modulates OSN synaptic output) ^46, 47^ can cause corresponding changes in the generalization of fear across odors ^10^.

**Figure 5.**
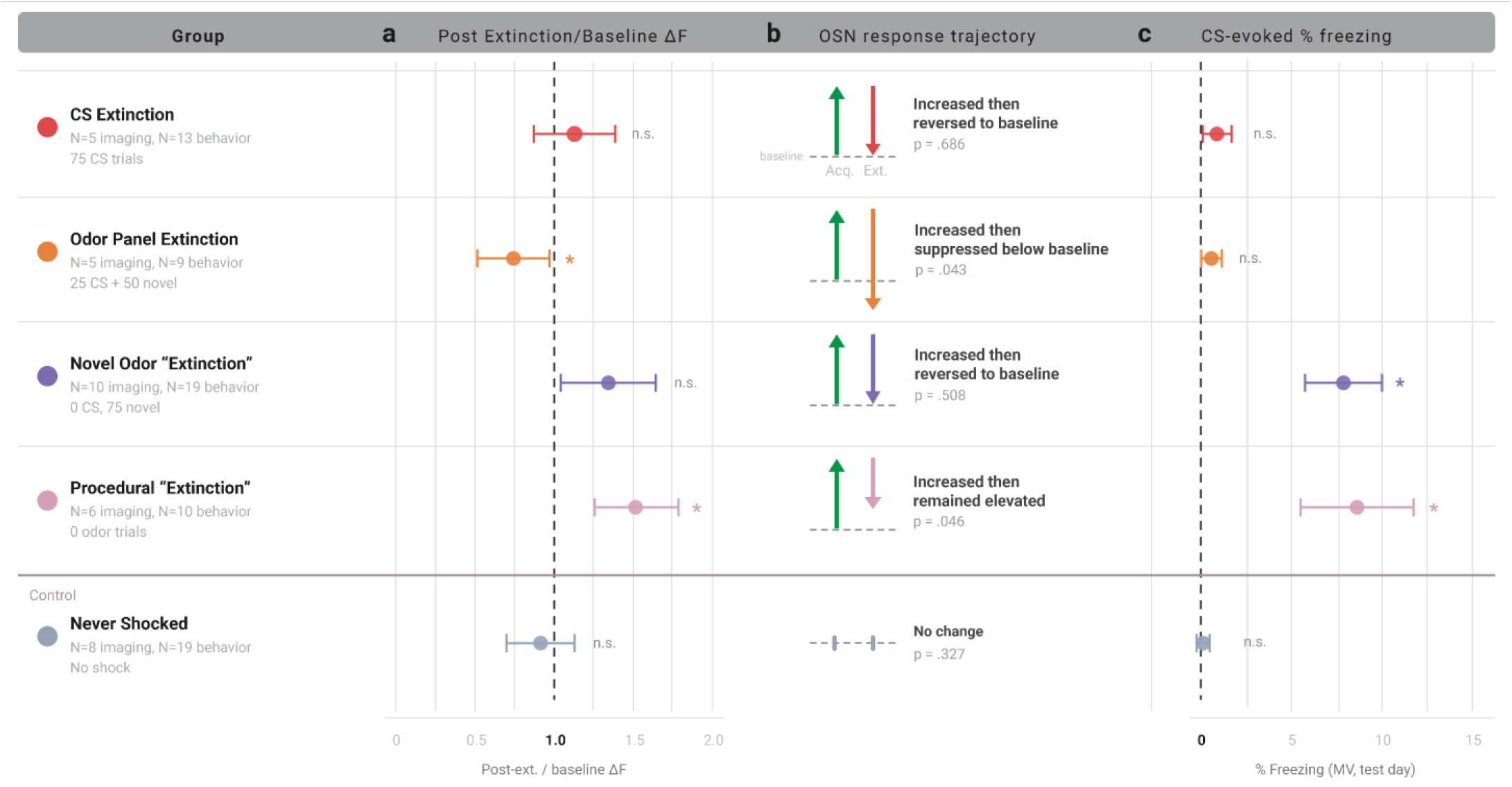
Summary of effects across extinction paradigms on CS odors. The CS Extinction paradigm (top row) reduced freezing to zero while returning odor-evoked OSN output to baseline. The Odor Panel Extinction paradigm (second row) reduced freezing to zero while reducing odor-evoked OSN output significantly below baseline despite fewer CS presentations than CS extinction. Mice undergoing the Novel Odor “Extinction” paradigm (third row) retained some freezing to the CS, along with a moderate though non-significant elevation in OSN output. Mice that underwent the Procedural “Extinction” control paradigm (fourth row), in which no explicit odors were presented during extinction training, also retained both some freezing to the CS and significant elevation in CS-evoked OSN output. Mice in the Never Shocked control group, which were never shocked but received the same odor presentations as mice in the CS Extinction group, exhibited no freezing and no significant change in OSN output. Graphs indicate mean ± SEM for the MV odor.

## Discussion

These experiments demonstrate multiple new findings about how olfactory fear conditioning affects the early olfactory representation of the threat-predictive odor, including a) that a single day of odor-cued fear conditioning increases the OSN synaptic output evoked by the threat-predictive odor (Fig. 1f**&i**), b) that this effect can survive 5 days of contextual extinction (Fig. 1h**&k**), and c) that conventional extinction training with the CS odor reverses both the behavioral fear evoked by that odor and the corresponding facilitation of the OSN response to that odor (Fig. 1f**&i**). However, they also provide new insight into the effects of olfactory fear conditioning on other odors that do not explicitly predict the shock, including a) that behavioral generalization of fear to novel odors was accompanied by corresponding increases in the OSN response to those odors (Fig. 2a**-e**), b) that these increases occurred even in OSN populations that were not responsive to the threat-predictive odor itself (Fig. 2f), and c) that the overall effect of these increases was to make the representations of odors substantially more different in an absolute sense but somewhat more similar in relative pattern independent of response amplitude (Fig. 4b, d, e). Finally, these data demonstrate the effects of new extinction training paradigms exploring the “generalization” of extinction learning to odors the mouse had previously generalized to, showing a) that extinction with a panel of odors including the CS extinguished conditioned fear more rapidly (compared to the same number of CS presentations) and more broadly across odors than conditioning with the CS alone (Fig. 3g), b) that odor panel extinction more than reversed the conditioning-induced enhancements of OSN responses to odors, leaving them smaller than the pre-conditioning baseline (Fig. 3c), and c) extinction-like training with a single novel odor was less effective than both conventional extinction training and odor panel extinction training at reducing conditioned fear, but did partially generalize to the CS, including partially reversing the facilitation of CS-evoked OSN output (Fig. 3d **& h**).

The reversal of the facilitation of CS-evoked OSN output by conventional extinction training with the CS is consistent with a previous demonstration that CS-specific glomerular structural enhancements are reversed by extinction of olfactory fear learning over a multi-week timescale ^33^. However, this result is in some ways surprising because extinction learning famously preserves some trace of the original acquisition learning, as revealed by phenomena like rapid reacquisition, spontaneous recovery, reinstatement, and renewal ^35, 48^. If the purpose of sensory changes during fear learning is indeed to enhance sensitivity to an ecologically critical stimulus ^1, 11^, this heightened sensitivity or salience for the previously threat-predictive stimulus would have been an appealing mechanism for some of these post-extinction “savings” effects. Additional experiments employing reacquisition training or informed by new mechanistic insights into the cellular and molecular basis of the fear-induced OSN plasticity may shed light onto the nature of any post-extinction memory trace in the OSNs.

The fear conditioning paradigm employed in these experiments was explicitly designed to promote generalization across odors, principally by using a single odor, limited number of trials, and a strong shock ^20^. As expected, mice exhibited clear behavioral generalization to all tested odorants, including odors they had never experienced previously while awake. This generalization of behaviorally expressed fear was paralleled by significant facilitation of the OSN response to all tested odors, even the novel ones. This finding is consistent with similar effects noted throughout the early olfactory processing circuit, including periglomerular cells ^20^, mitral cells ^7^, and anterior piriform cortex ^21^. Given the large size of the fear conditioning effect at the level of the OSNs, our data suggest that the corresponding effects at downstream neurons are principally reflecting their increased input from OSNs following learning.

A key advantage of the olfactory system for exploring the mechanism of generalization is that because of the anatomical mapping of odor receptor onto olfactory bulb glomerulus it is possible to independently observe the neural representation of each chemical feature of the odorant molecule. Similar odors activate overlapping sets of receptors, enabling the direct comparison of effects on glomerular populations of OSNs that respond to both the CS odor and another odor in the panel (i.e. that represent shared stimulus “features” between the CS and the odor the mouse generalizes to) and effects on glomerular populations of OSNs that respond to only to a novel odor and not to the CS (i.e. that represent stimulus “features” absent from the CS). In our data the facilitation of odor-evoked OSN neurotransmitter release was observed in *both* types of glomeruli, demonstrating that the facilitated OSN output evoked by novel odors is not due to any overlap in chemical features with the CS. This mirrors the overall pattern of behavioral and neural generalization across odors, where mice were similarly afraid of all odors and OSNs were similarly facilitated across odors, regardless of their chemical or perceptual similarities.

If not based on shared chemical features, how does the overall response pattern of OSN output (i.e. the configural representation of the odor) change after fear learning? We explored this by representing the response patterns across glomeruli in an N-dimensional space where each dimension is defined by the OSN output into a glomerulus (corresponding to the response to a chemical feature of the odorant). The relative size of the responses among glomeruli thus determines the angle (direction of travel) of the neural representation in this space during odor presentation, while the amplitude of the responses determines the amplitude (distance traveled) of the neural representation ^49^. We observed that fear learning increased the absolute (Euclidean) distances between neural representations of the CS and each novel odor, reflecting the overall larger neural responses after fear conditioning (and consistent with previous reports of improved fine discrimination between similar odors and improved odor sensitivity ^14, 15, 17^). However, the changes in cosine difference after conditioning suggest that two odor representations became more similar to the CS after fear conditioning. This similarity to the CS could reflect an increased perceptual similarity between them or alternatively the perceptual quality might remain unchanged while the neural representations directly incorporate information about an odor’s ecological significance or priority.

The three extinction paradigms tested here produced a graded set of outcomes at both the behavioral and neurophysiological levels. Conventional CS extinction fully reversed the facilitation of CS-evoked OSN output and eliminated CS-evoked freezing, but left residual facilitation for some novel odors and residual freezing to 2-Hex, the odor most dissimilar to the CS. Novel odor “extinction” reversed OSN facilitation broadly across odors, including those not presented during extinction, but left intact significant freezing to the CS itself. Odor panel extinction produced the most complete reversal: all tested odors returned to or below pre-conditioning baseline in both OSN output and conditioned freezing, despite including only 15 CS presentations compared to 75 in the conventional paradigm. The Procedural “Extinction” control, in which mice underwent the extinction procedure without odor presentations, produced only partial reversal of both measures. This ordering of outcomes was consistent across neurophysiology and behavior (Fig. 5), with the paradigms that left the least residual OSN facilitation also producing the least residual freezing, and vice versa.

The effects of extinction training on sensory processing have particular translational significance because exposure therapy delivered in the clinic following a trauma is essentially extinction training. The notable efficacy of the odor panel paradigm in reversing both the behavioral changes and neurophysiological plasticity induced by odor-shock pairings suggests that exposure to a range of stimuli ranging in similarity to the original CS might be equally if not more effective to reverse the generalization of behaviorally expressed and neural expressed fear. These results also raise the important idea that conventional fear conditioning and extinction might be a poor model of anxiety disorders because the subject is correct to be afraid and slow to reverse course. Inappropriate generalization is a superior model of disordered fear, and the ability to narrow that fear to just the stimuli that are most appropriate has potential implications for clinical therapies. Taken together, these experiments demonstrate the fundamental relationship between learning, sensory processing, and perceptual plasticity and supports our previous findings that disruption of neurosensory plasticity may be part of the etiology or maintenance of anxiety ^17^.

The initial finding that fear conditioning with a single odor on a single day induces large increases in the OSN synaptic output evoked by that odor is new but not unexpected. It is consistent with previous effects observed using the same paradigm in periglomerular neurons immediately downstream of the OSNs ^20^, suggesting that the increased PG cell activity after fear learning indeed reflects increases in their synaptic input from the periphery. It is also consistent with previous work in OSNs using three days of discriminative olfactory fear conditioning ^11, 13^ and using extended periods of single odor conditioning or odor-drug pairing ^9^. An overnight delay between conditioning and testing is not long enough to develop entirely new OSNs responsive to the CS odor in the olfactory epithelium, ruling out anatomical explanations ^50^. Alternatively, OSN synaptic output is strongly modulated by GABA_B_ receptors on their presynaptic terminals ^46, 47, 51^, which is reduced after odor-cued fear learning for CS-responsive OSNs ^10, 52^. GABAergic signaling in the glomeruli altered by fear learning ^20^ and is modulated by amygdala-driven noradrenergic projections from the locus coeruleus to the olfactory bulb ^53^ and thus makes a strong candidate mechanism for rapid, fear learning-induced neuroplasticity in OSN terminals.

The fear conditioning-induced facilitation of OSN neurotransmitter release evoked by novel odors is particularly notable because the mice were anesthetized during imaging sessions. The olfactory bulb is richly innervated by centrifugal projections from structures like the locus coeruleus, whose release of norepinephrine into the OB is modulated by amygdala output ^53^, the basal forebrain’s rich cholinergic inputs to periglomerular regions ^54^, massive reciprocal connections with piriform cortex ^55–57^, and hippocampus ^58, 59^. The presentation of a fear-evoking odor in an awake animal evokes substantial changes in all of these circuits along with large autonomic changes in breathing and heart rate that could collectively have drastic effects on stimulus processing in the olfactory bulb and all over the brain. Such changes might be characterized as “retrieval-mediated” or even considered part of the conditioned response. However, in anesthetized mice no such autonomic responses are observed, including no change in respiration ^11^, and olfactory processing changes are readily observable as early as the rising phase of the first inhalation of odor ^20^. We thus expect that the relevant neuroplasticity happens during learning and that the fear-induced plasticity is already encoded locally in the earliest olfactory circuitry prior to odor presentation.

The OSNs have direct knowledge of odors in the environment and likely have indirect knowledge of footshocks (via endocrine or neuromodulatory signaling), so the initial learning of the CS odor-shock contingency could occur via covariance-based plasticity (e.g. Hebbian synapses). However, the present work demonstrates the generalization of learning effects to OSN populations that are not responsive to the shock-predictive odor, the generalization of extinction from novel odors to the CS odor, and the reversal of prior learning effects by the *absence* of a footshock. These phenomena are not readily implemented by a local covariance rule in the OSNs and suggest that even the earliest parts of the olfactory system participate in a larger network of cognitive function.

## Acknowledgements

This work was funded by R01 MH101293 from the National Institute of Mental Health and the National Institute on Deafness and other Communication Disorders. We thank Walter Shotwell, Adam Garcia, and Jayanne Pierre for technical assistance and Kasia Bieszczad for helpful suggestions on experimental manipulations.

## Conflicts of Interest

The authors declare no conflicts of interest.

## Notes

### Competing Interest Statement

The authors have declared no competing interest.

